# Thermal mismatches explain how climate change and infectious disease drove widespread amphibian extinctions

**DOI:** 10.1101/164814

**Authors:** Jeremy M. Cohen, David J. Civitello, Matthew D. Venesky, Taegan A. McMahon, Jason R. Rohr

## Abstract

Global temperatures and infectious disease outbreaks are simultaneously increasing, but linking climate change and infectious disease to modern extinctions remains difficult. The *thermal mismatch hypothesis* predicts that hosts should be vulnerable to disease at temperatures where the performance gap between themselves and parasites is greatest. This framework could be used to identify species at risk from a combination of climate change and disease because it suggests that extinctions should occur when climatic conditions shift from historical baselines. We conducted laboratory experiments and analyses of recent extinctions in the amphibian genus *Atelopus* to show that species from the coldest environments experienced the greatest disease susceptibility and extinction risk when temperatures rapidly warmed, confirming predictions of the *thermal mismatch hypothesis*. Our work provides evidence that a modern mass extinction was likely driven by an interaction between climate change and infectious disease.

---

Global climate change and emerging infectious diseases represent two of the most formidable ecological challenges in modern times, but controversy exists over whether they are causally linked (*1, 2*). Climatic conditions often directly influence disease outbreaks (*3*), and many predictive models and experiments have revealed that climate change and infectious diseases can independently drive current and future declines in biodiversity (*4, 5*). However, there is surprisingly little concrete evidence that interactions between climate change and infectious disease are causing widespread biodiversity losses (*1, 6*), possibly because of a lack of theoretical frameworks, supported by a combination of experiments and field data, that can relate climatic factors to host-parasite interactions to account for shifts in biodiversity (*1*). Such frameworks would be valuable in establishing causal links between climate change and extinctions mediated by disease.

A recently proposed hypothesis, the *thermal mismatch hypothesis* (*7*), suggests that infectious disease outbreaks are likely to occur at temperatures where the performance gap between pathogens and their hosts is greatest. Because parasites generally have broader thermal performance breadths than hosts (*8, 9*), and both hosts and parasites might be locally adapted to climatic conditions in their ranges and limited by extreme conditions, the hypothesis posits that hosts adapted to cooler climates should be especially susceptible to disease under unusually warm conditions, and vice versa (Fig. 1). Importantly, the predictions of the thermal mismatch hypothesis are robust to relaxing several of its assumptions, such as local adaptation of host and parasite and the degree and direction of the skew of the performance curves (Fig. S1). Therefore, the *thermal mismatch hypothesis* provides a framework to predict which species might be most likely to experience disease-driven declines under warming, and thus might be able to explain patterns in species declines associated with climate-related outbreaks of emerging infectious diseases.

**Fig. 1.**
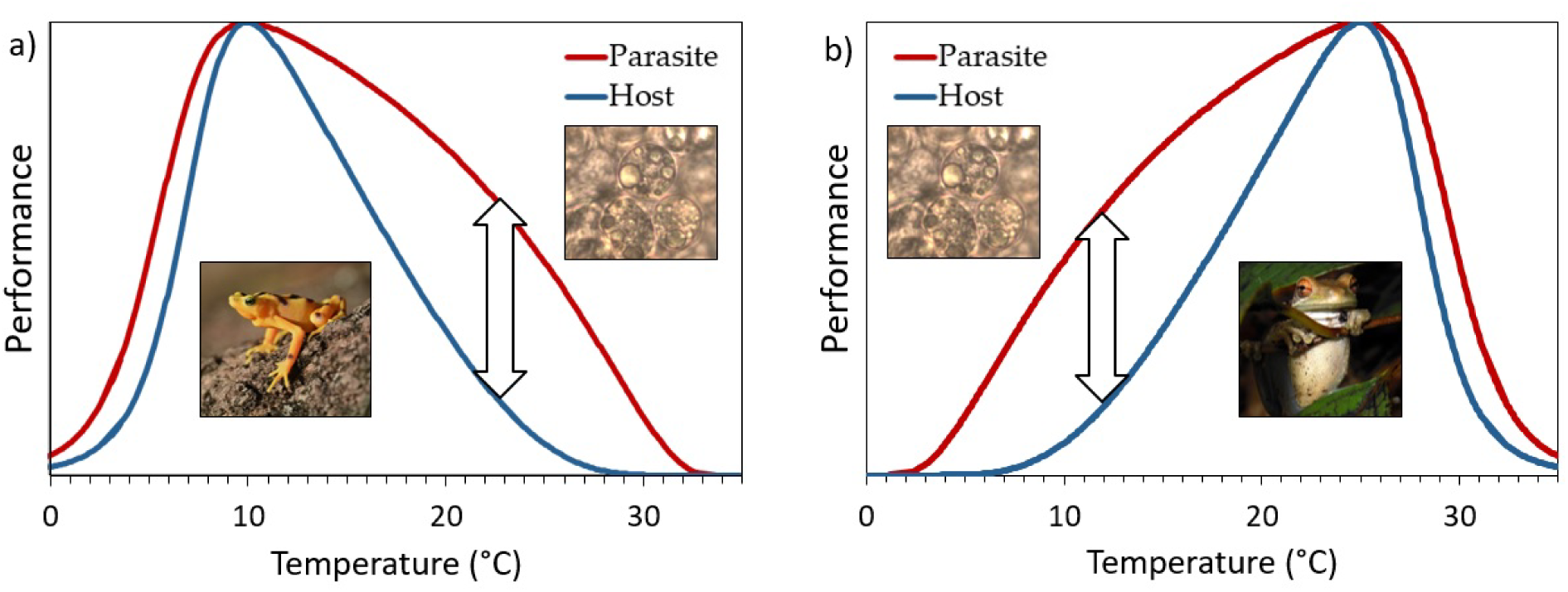
Conceptual figure describing the *thermal mismatch hypothesis*. In isolation, small organisms, such as parasites (red lines), generally have broader thermal performance curves than larger organisms, such as hosts (blue lines). Parasite growth on hosts is likely to occur at temperatures where parasite most outperforms its host (bidirectional arrows), and not necessarily at the temperature at which a parasite performs best in isolation, providing a hypothesis for the thermal performance curve of a parasite growing on the host. For interacting cold-adapted hosts and parasites **(a)**, parasite growth should be maximized at relatively warm temperatures, whereas for interacting warm-adapted hosts and parasites **(b)**, parasite growth is predicted to be maximized at relatively cool temperatures.

Although recent experimental and field evidence support the predictions of the *thermal mismatch hypothesis* (*7*), the hypothesis has not yet been applied to predict widespread host declines associated with climate change and infectious disease. In perhaps the most comprehensive spatiotemporal dataset describing a modern mass extinction, La Marca *et al*. (*10*) provide dates of approximately 60 extinctions in the amphibian genus *Atelopus* putatively caused by the chytrid fungus *Batrachochytrium dendrobatidis* (*Bd*), a pathogen that infects the epidermal layer of adult amphibians and is implicated in worldwide amphibian declines (*11; Table S1*). These *Atelopus* declines have been linked to climate change (*12, 13*), occurring in years with warm or highly variable temperatures (*12, 13*) despite *Bd* growing best in culture under cool or moderate conditions (*7*). *Bd* is sensitive to environmental conditions (*14*), can be locally adapted (*15*), and has a broad thermal breadth (*16*), fulfilling the assumptions of the *thermal mismatch hypothesis*. Thus, this *Atelopus* extinction dataset provides a unique opportunity to examine whether the *thermal mismatch hypothesis* can accurately predict the timing and location of disease-driven extinctions and to compete the *thermal mismatch hypothesis* against alternative hypotheses for these climate-related declines. Although there have been previous analyses associating climate change and *Atelopus* spp. extinctions, they relied exclusively on correlations based on broad-scale, regional climate data instead of data from individual species’ ranges (*12, 13; Supplement*) and thus they failed to account for spatial heterogeneity in climate change and host adaptations to local climates. Therefore, previous analyses could not determine whether *Atelopus* spp. declines were likely caused by climate change alone or an interaction between climate change and disease. Thus, we do not yet have clear, causal evidence in this system, or any system that we are aware of, that climate change caused extinctions by facilitating infectious disease (but see *17*).

Here, we take a hypothetico-deductive approach, linking a theoretical framework, laboratory experiments, and analyses of field data to examine the relationships among extinctions, climate change, and emerging infectious disease. We simultaneously tested six hypotheses or predictors for the climate-related *Atelopus* spp. extinctions: 1) a null model, 2) pathogen alone: temperature-dependent growth of *Bd* in culture, 3) temperature variability alone: annual month-to-month variability in temperature, 4) mean climate alone: annual mean temperature, 5) climate change alone: the 5-year slope of mean temperature, and 6) the interaction between mean historical climate and climate change: because the *thermal mismatch hypothesis* predicts that the effect of climate change depends on whether the host is cool or warm adapted, which in turn drives the differential performance of host and pathogen.

Given that previous climate change analyses of the *Atelopus* dataset relied on correlating extinctions with regional climate data (*12, 13*), we first verified that climate change was indeed associated with these extinctions based on data from individual species’ ranges. In the geographic ranges of species that went extinct, mean temperatures in the five years leading up to extinction increased ∽2.5 times faster than they increased in the ranges of species that remained extant (matched pairs test: *F*_*1,45*_=7.73, *p*<0.01; see Methods; Fig. S2; Table S2) (see *18 for a similar approach using two species*). Hence, soon-to-be extinct species were indeed experiencing conditions that were both unusually warm for them and warmer than those experienced by species that remained extant, consistent with the hypothesis that climate change played a role in *Atelopus* spp. declines.

Next, we set out to parameterize our statistical model by conducting laboratory experiments to evaluate the impacts of both mean temperature and temperature variability on *Atelopus* spp. mortality risk. First, we conducted a *Temperature Gradient Experiment* in which we exposed *Atelopus zeteki*, which we consider to be cold-adapted for a variety of reasons (see Supplement), to *Bd* in replicated temperature-controlled incubators (*19; Fig. S2*) across a naturally relevant temperature gradient (14°, 18°, 22°, 26°, and 28°C) while simultaneously maintaining unexposed frogs and growing *Bd* in liquid cultures in the same incubators. In this experiment, the temperature gradient did not affect *A. zeteki* mortality in the absence of *Bd* (cox proportional-hazards model: *X*^*2*^ = 0.54, *p* = 0.46), but mortality increased significantly with temperature when *A. zeteki* was exposed to *Bd* (*Bd x* temperature: *X*^*2*^ = 4.41, *p* = 0.036). In fact, within a week of exposure to *Bd*, frogs at 26° and 28°C experienced 69% and 78% mortality, respectively, suggesting a temperature-dependent cost of exposure to *Bd* (see Supplement), whereas only one *Bd*-exposed animal died at the two coldest temperatures within a week of exposure (6% mortality) and only four *Bd*-negative animals died throughout the experiment (20% mortality; Fig. 2; Fig. S4). Similarly, *Bd* growth rates on frogs increased with temperature (Fig. 2a). In contrast, temperature-dependent *Bd* growth in culture closely followed previously reported patterns with growth rates increasing as temperature increased until 18.0°C (optimum) and then decreasing thereafter with little growth above 26°C (*20; Fig. 2a, Fig. S5*). These results demonstrate that patterns of temperature-dependent *Bd* performance in culture and on hosts differ sharply, a result consistent with the *thermal mismatch hypothesis*, which predicts that parasites should have maximum growth on host at temperatures where they most outperform the host rather than at temperatures where the parasite has the greatest absolute performance in culture. The striking monotonic positive association between temperature and both *Bd* growth on frogs and *Bd*-induced host mortality contradict a common assumption that *Bd* outbreaks only occur at cool or moderate temperatures (*14, 21*). Importantly, although we only tested one *Atelopus* species in this experiment, the observed patterns are likely generalizable to other *Atelopus* spp., because a global analysis of *Bd* prevalence in 15,410 individuals from 598 amphibian populations and 1,399 species revealed that cold-and warm-adapted amphibians generally have peak *Bd* prevalences during warm and cold spells, respectively (*7; Fig. S6*).

**Fig. 2.**
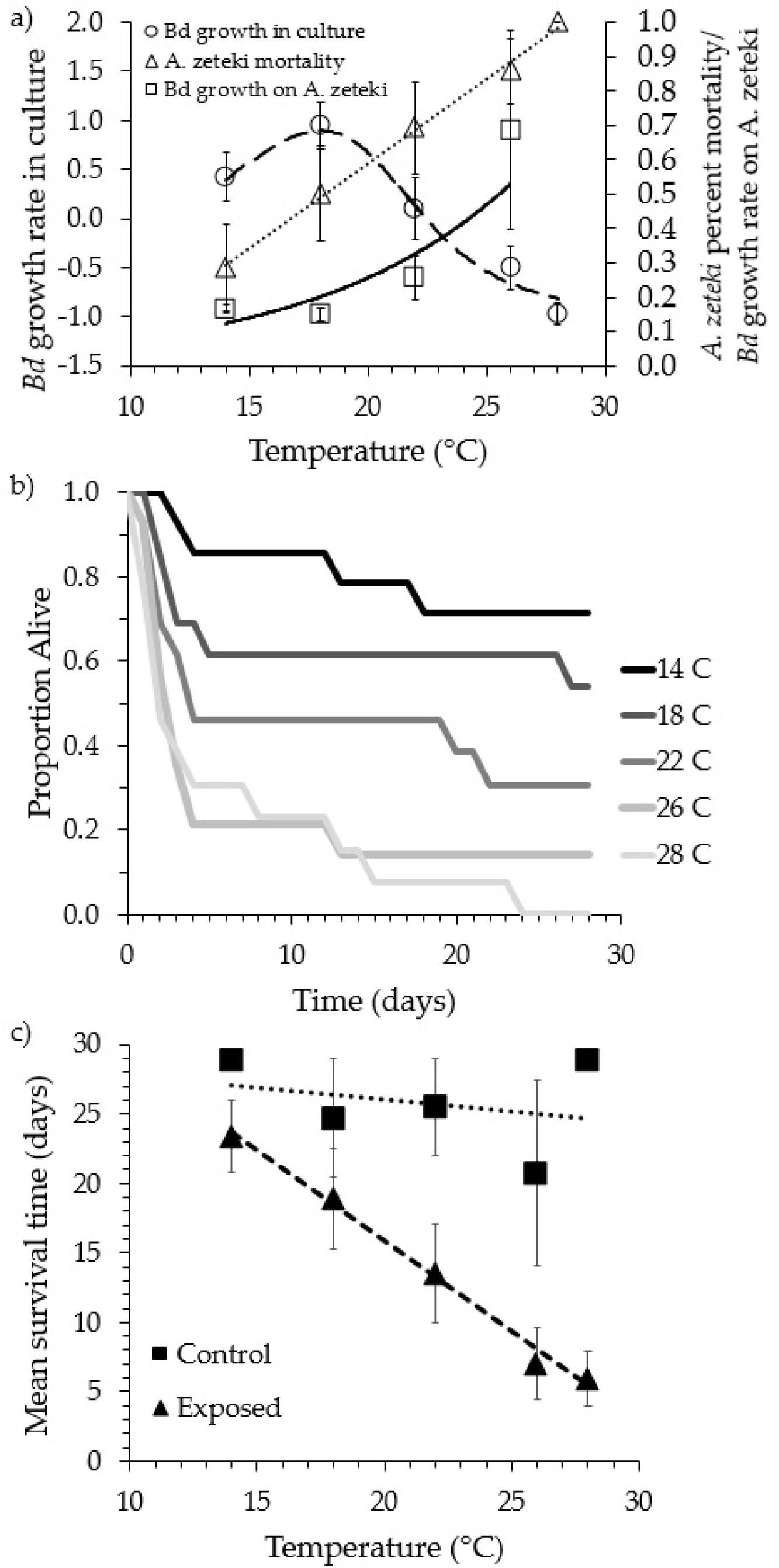
Temperature-dependent patterns of *Batrachochytrium dendrobatidis* (*Bd*) growth and *Atelopus zeteki* mortality. **(a)** *A. zeteki* experienced high mortality (triangles, dotted line) and high *Bd* growth (squares, solid line) at warm temperatures after *Bd* exposure, even though *Bd* growth rates in culture (circles, dashed line) were low at these temperatures. We could not measure *Bd* growth rates on *A. zeteki* at 28 °C because very few animals survived long enough to be tested multiple times. **(b)** Survival plot for *A. zeteki* exposed to *Bd* across two temporal blocks. **(c)** Mean survival time for *A. zeteki* at each of five temperatures when exposed to *Bd* (triangles; both temporal blocks) or not exposed (squares; 2^nd^ temporal block). Temperature and *Bd* exposure interacted to induce high mortality in *A. zeteki* (*X*^*2*^ = 4.41, *p* = 0.036). Animals surviving the experiment are conservatively assumed to have died on day 29 for these figures only but were censored in the survival analysis. Error bars represent SEMs in all panels.

Second, we conducted a *Temperature Shift Experiment* in which we exposed *A. zeteki* to *Bd* at either constant or variable temperatures to evaluate how temperature variability affects host susceptibility. *A. zeteki* were exposed to *Bd* at 14°, 17°, 23°, or 26°C immediately following either two weeks of acclimation to these temperatures or two weeks of acclimation to 20°C, so that all the hosts experienced either constant or shifted temperatures before *Bd* exposure, respectively. As with the previous experiment, *Bd*-induced mortality increased with temperature (cox-proportional hazards model and ANOVA: *X*^*2*^=4.08, *p*<0.05; Table S3). At the same *Bd* exposure temperatures, frogs that experienced temperature shifts had higher *Bd* loads than those that did not experience shifts (ANOVA: *F*_*1,34*_=8.78, *p* = 0.005), consistent with the findings of previous studies (*19, 22*). However, we did not observe any significant effect of the temperature shift treatment on mortality (Shift treatment: *X*^*2*^=0.84, *p*=0.36; Shift *x* temperature: *X*^*2*^=1.03, *p*=0.31), and the temperature gradient accounted for >6 times the variance in *Bd*-induced mortality as temperature variability (Table S4).

Given the results of our two laboratory experiments, we hypothesized that *Bd* growth in culture, temperature variability, and mean temperature alone would be poor predictors of *Atelopus* extinctions in the wild relative to the *thermal mismatch hypothesis*, which posits that as temperature increases, disease and extinction risk should be most pronounced among *Atelopus* spp. from cooler regions because they should experience a larger performance gap relative to *Bd* than species from warmer regions (Fig. 1). This prediction of the *thermal mismatch hypothesis* would manifest as a statistical interaction between the temperature to which a species is adapted (50-year mean temperature in a species’ geographic range) and the level of climate change it has experienced because cold-adapted species should experience disease *-* associated declines when temperatures increase, whereas warm-adapted species should not. To test these hypotheses, we utilized a time-dependent cox-proportional hazards survival model (*23, see Methods*) that concurrently evaluated the following predictors of the occurrence and timing of extinctions: *Bd* growth in culture, temperature variability, mean temperature, climate change, and the *thermal mismatch hypothesis* (see Methods). Given that extinction probabilities have repeatedly been shown to be negatively dependent on geographic range size (*24*), range size was included as a crossed factor with each predictor in our model. The model also controlled for two precipitation variables and altitude, which have been associated with *Atelopus* spp. extinction probabilities (*12*).

Consistent with our experiments, *Atelopus* spp. extinction risk was not significantly explained by interactions between geographic range size and *Bd* growth in culture or temperature variability but was significantly explained by the *thermal mismatch hypothesis* (Table 1). Species with large range sizes rarely experienced extinctions and thus were not strongly impacted by climate or disease. In contrast, species with smaller range sizes showed extinction patterns consistent with the *thermal mismatch hypothesis*. Increasing temperatures associated with climate change (positive slope five years before extinction) were positively associated with the occurrence and timing of the extinction of cold-adapted *Atelopus* spp. (Fig. 3a,c), whereas climate change did not predict the occurrence and timing of declines of warm-adapted species (Range size *x* temperature shift *x* 40-year mean temperature; *β*=11.5, df=22, *p*=0.02, Table 1, Fig. 3b,d). In fact, in the absence of any climate change, warm-adapted species were more likely to experience extinctions in cool rather than warm years (Fig. 3b,d), also consistent with the *thermal mismatch hypothesis* (Fig. 1). The model testing the *thermal mismatch hypothesis* explained about 2.5 times more of the variance in extinctions than a model that did not contain the interaction (Nagelkerke’s pseudo-*R*^*2*^=0.466 and 0.189, respectively).

**Table 1.** Results of time-dependent cox-proportional hazards model predicting extinction with a four-way interaction between log-transformed range size, long-term mean temperature (40yr.meantemp), annual mean temperature (meantemp), and recent temperature shift (tempchange) across both extinct and extant *Atelopus* spp. Mortality probability based on *Bd* growth in culture (culturemortprob), log-transformed altitude (logaltitude) and a measure of temperature variability (log-transformed AVMD, absolute value of monthly difference in temperature) were also included. Bolded lines represent tests of the *thermal mismatch hypothesis*.

|  | Coefficient | SE | Robust<br>SE | z | p |
| --- | --- | --- | --- | --- | --- |
| Rangesize | -23.500 | 72.300 | 51.500 | -0.460 | 0.64 |
| Logaltitude | -0.124 | 0.283 | 0.147 | -0.840 | 0.39 |
| Culturemortprob (pathogen only) | -1.130 | 1.480 | 1.220 | -0.930 | 0.35 |
| Total precipitation | <0.001 | <0.001 | <0.001 | 0.840 | 0.40 |
| Frequency of wet days | <0.001 | <0.001 | <0.001 | -0.280 | 0.77 |
| Log(AVMD) (temp. variability) | 0.913 | 0.756 | 0.719 | 1.270 | 0.20 |
| Tempchange (climate change only) | 11.300 | 35.600 | 37.600 | 0.300 | 0.76 |
| 40yr.meantemp (cold or warm adapted) | 1.310 | 0.901 | 0.759 | 1.730 | 0.08 |
| Meantemp (mean temp. only) | 0.564 | 0.972 | 0.756 | 0.750 | 0.45 |
| Rangesize:Culturemortprob | -4.070 | 9.130 | 6.690 | -0.610 | 0.54 |
| Rangesize:log(AVMD) | -1.410 | 2.030 | 1.810 | -0.780 | 0.43 |
| Rangesize:Tempchange | -160.000 | 155.000 | 82.800 | -1.930 | 0.05 |
| Rangesize:40yr.meantemp | 1.460 | 4.380 | 2.890 | 0.510 | 0.61 |
| Tempchange:40yr.meantemp | -0.080 | 2.650 | 2.710 | -0.030 | 0.97 |
| Rangesize:meantemp | 3.130 | 5.610 | 4.040 | 0.770 | 0.43 |
| Tempchange:meantemp | 0.420 | 2.380 | 2.200 | 0.190 | 0.84 |
| 40yr.meantemp:meantemp | -0.051 | 0.040 | 0.033 | -1.580 | 0.11 |
| **Rangesize:Tempchange:40yr.meantemp** | **11.500** | **9.800** | **5.060** | **2.280** | **0.02** |
| Rangesize:Tempchange:meantemp | 5.680 | 9.480 | 6.130 | 0.930 | 0.35 |
| Rangesize:40yr.meantemp:meantemp | -0.146 | 0.280 | 0.195 | -0.750 | 0.45 |
| Tempchange:40yr.meantemp:meantemp | -0.028 | 0.102 | 0.111 | -0.250 | 0.80 |
| **Rangesize:Tempchange:40yr.meantemp:meantemp** | **-0.464** | **0.467** | **0.224** | **-2.070** | **0.03** |

**Fig. 3.**
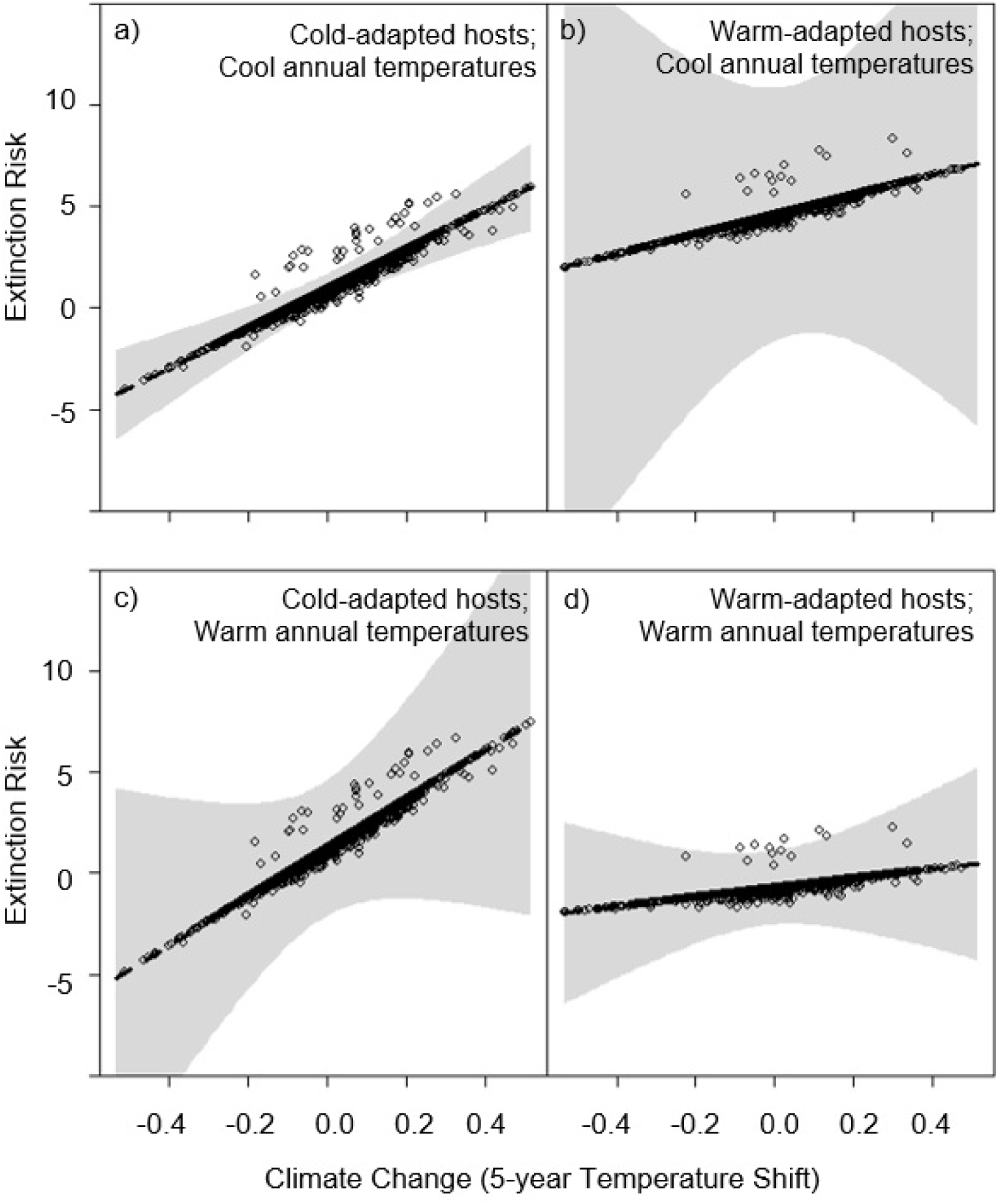
Partial residual plot displaying the effects of climate change and annual mean temperature on the extinction risk of cold-and warm-adapted *Atelopus* spp. The partial residuals are from the time-dependent cox proportional-hazards model shown in Table 1 and display the significant three-way interaction among 5-year slopes in mean temperature-by-40-year mean temperature-by-annual mean temperature. Points represent individual years for each species and gray shading shows associated 95% confidence bands. The model suggests that species from typically cooler climates (**a,c**) were at greater risk of extinction (log-odds risk ratio; y-axis) after experiencing climate change (warming, or positive 5-year slope in mean temperature; x-axis) than species from warmer climates (**b,d**; breaks based on 20^th^ and 80^th^ percentiles). This pattern was consistent whether the annual mean temperature in a given year was relatively cool (**a,b**) or warm (**c,d**; breaks based on 20^th^ and 80^th^ percentiles), although species from warmer climates may be at greater risk when conditions are cooler (**b**).

To gather further support for the notion that an interaction between climate change and *Bd* drove *Atelopus* extinctions, we compared the magnitude of climate change and extinctions experienced by genus *Atelopus*, which is believed to have been widely exposed to *Bd* and is found in a region of South America where *Bd* has been detected as early 1894 (*25*), to amphibians in Madagascar and Scandinavia, regions historically considered to be free of *Bd* (*26-28*). Compared to *Atelopus* spp., amphibian species in Madagascar and Scandinavia experienced similar and more climate change between 1950 and 2004, respectively (T-test: *T*=0.118, *p*=0.906; *T* =-5.59, *p*<0.0001; Fig. S7). However, unlike genus *Atelopus*, there were no amphibian extinctions in these areas during this time (*29*). This suggests that, in the absence of *Bd*, the same level of climate change experienced in Latin America was insufficient to cause amphibian extinctions in Madagascar and Scandinavia. Although there are major differences in taxonomy and life history between these groups of amphibians, as well as differences in the potential stressors they experienced, these results are consistent with the *thermal mismatch hypothesis* and our laboratory findings, which suggest that *Atelopus* declines were caused by an interaction between climate change and *Bd* rather than either stressor alone. However, we caution that these analyses assume a consistent response to climate change across regions, which may not be realistic given that a degree change in temperature has a greater metabolic impact on tropical species than it does on temperate species (*30*).

Our experiments and analyses of field data together suggest that *Atelopus* spp. from cooler environments are more vulnerable to mortality from chytridiomycosis under warmer conditions than those from warmer environments and thus, climate change poses a greater threat to cold-adapted *Atelopus* spp. Therefore, the *thermal mismatch hypothesis* was a useful framework for predicting which species were most likely to be impacted by an interaction between climate change and infectious disease outbreaks in this system. However, the generality of the *thermal mismatch hypothesis* is unknown, as it has only been evaluated in systems with ectothermic hosts and directly transmitted pathogens, which may be especially sensitive to environmental conditions. Further large-scale analyses of disease datasets are needed to test how well the *thermal mismatch hypothesis* applies across host-parasite systems that vary in host thermal biology or mode of transmission.

As global temperatures and infectious disease outbreaks have increased, these two crises have been repeatedly correlated by researchers to explain species declines and extinctions. However, evidence that they interact to cause declines has been elusive, possibly because researchers have tried to simplistically correlate increases in temperature with infectious disease, rather than looking for more nuanced patterns that depend on the host-parasite interaction (*1*). Here, we apply the *thermal mismatch hypothesis*, a framework that can relate environmental temperature to disease patterns while accounting for host-level variation in adaptation to climate to predict which host species are most vulnerable to infectious disease with global warming. By combining experiments with field patterns to examine how mean temperature and temperature variability impact susceptibility to *Bd* in the amphibian genus *Atelopus*, we provide the first evidence that one of the greatest modern day mass extinctions was likely driven by an interaction between climate change and infectious disease.

## Acknowledgments

Thanks to the Maryland Zoo for providing us with the *A. zeteki* used in the experiments. We thank N. Argento, K. Ebener, H. Folse, C. Gionet, T. James, C. Koshy, C. Malave, L. Martinez-Rodriguez, C. Steffan, S. Rubano, and K. Vazquez for their assistance in all aspects of the lab work and animal maintenance. In addition, we thank all members of the Rohr lab for their helpful comments on the manuscript. Funds were provided by grants to J.R.R. from the National Science Foundation (EF-1241889), the National Institutes of Health (R01GM109499, R01TW010286-01), the US Department of Agriculture (2009-35102-0543), and the US Environmental Protection Agency grant (CAREER 83518801).

## Author contributions

All authors contributed ideas, J.M.C. and M.D.V. wrote proposals to acquire animals, J.M.C., M.D.V. and T.A.M. conducted disease experiments, J.M.C. assembled climate database, J.M.C., D.J.C. and J.R.R. conducted statistical analyses, J.M.C. and J.R.R. wrote the paper, and all authors provided editorial advice.

